# Replacing the *SpCas9* HNH domain by deaminases generates compact base editors with an alternative targeting scope

**DOI:** 10.1101/2020.11.09.371237

**Authors:** Lukas Villiger, Lukas Schmidheini, Nicolas Mathis, Tanja Rothgangl, Kim Marquart, Gerald Schwank

## Abstract

Base editors are RNA-guided deaminases that enable site-specific nucleotide transitions. The targeting scope of these Cas-deaminase fusion proteins critically depends on the availability of a protospacer adjacent motif (PAM) at the target locus and is limited to a window within the CRISPR-Cas R-loop, where single stranded (ss)DNA is accessible to the deaminase. Here, we reason that the Cas9-HNH nuclease domain sterically constrains ssDNA accessibility, and demonstrate that omission of this domain expands the editing window. By exchanging the HNH nuclease domain with an adenosine deaminase we furthermore engineer adenine base editor variants (HNHx-ABE) with PAM-proximally shifted editing windows. This work expands the targeting scope of base editors, and provides base editor variants that are substantially smaller. It moreover informs of potential future directions in Cas9 protein engineering, where the HNH domain could be replaced by other enzymes that act on ssDNA.

## INTRODUCTION

CRISPR-Cas systems provide adaptive immunity to bacteria to defend them against foreign genetic elements. A small number of these systems have successfully been adopted for genome editing in mammalian cells, transforming biomedical research and therapeutics (1-3). A paradigmatic example is the *SpCas9* nuclease from *Streptococcus pyogenes.* Recruitment of *SpCas9* to a desired genomic locus via a single guide (sg)RNA allows facile and efficient genome editing by generating site-specific double stranded (ds)DNA breaks. Precise genome editing via such dsDNA breaks, however, relies on template DNA for homology directed repair (HDR) that is inefficient in most mammalian cell types and leads to repair via alternative, error-prone end-joining pathways that generate frameshift mutations and gene knockouts.

Recent modifications to the highly modular *Sp*Cas9 protein resulted in sophisticated genome editing technologies, such as base editors (BEs), which enable genome editing without dsDNA break formation. BEs allow precise conversion of A-to-G and C-to-T nucleobases or *vice versa* via nucleotide deamination independent of dsDNA breaks and HDR (4, 5), and thus have great potential for research- and therapeutic applications (6–8). They are mono- or heterodimers of ssDNA-specific deaminases fused N-terminally to catalytically impaired CRISPR class II effectors such as Cas9 or Cas12a (4, 5, 9). Upon binding to a protospacer element, Cas9 complexed with the sgRNA forms an R-loop, making the non-target DNA strand (the target DNA strand is bound to the sgRNA) accessible to ssDNA-specific deaminases. Notably, Cas9 and Cas12 proteins require PAM sites for DNA binding, restricting the targeting scope of base editors to nucleobases that lie within a defined proximity to these motifs.

To address the limited PAM compatibility, various base editors were generated by fusing deaminases to Cas variants with less restrictive PAM requirements (10–14). While these efforts considerably expanded the targeting scope of BEs, currently, no PAMless Cas9 and Cas12 variants have been developed, and several T>C and G>A disease-causing SNPs remain untargetable. Pursuing a different approach, recent work has led to base editor variants with slightly extended editing windows: Huang *et al.* fused deaminases to circularly permuted Cas9 proteins (15), and Wang *et al.* incorporated the cytidine deaminase rAPOBEC1 into the PAM-interacting (PI) domain of *Sp*Cas9 (16). Similarly, incorporation of laboratory evolved TadA7.10 (4) into the PI domain led to a PAM-proximal extension of the editing window of adenine base editors (ABEs) (ABEmax PI - Fig 1a and b).

**Fig 1.**
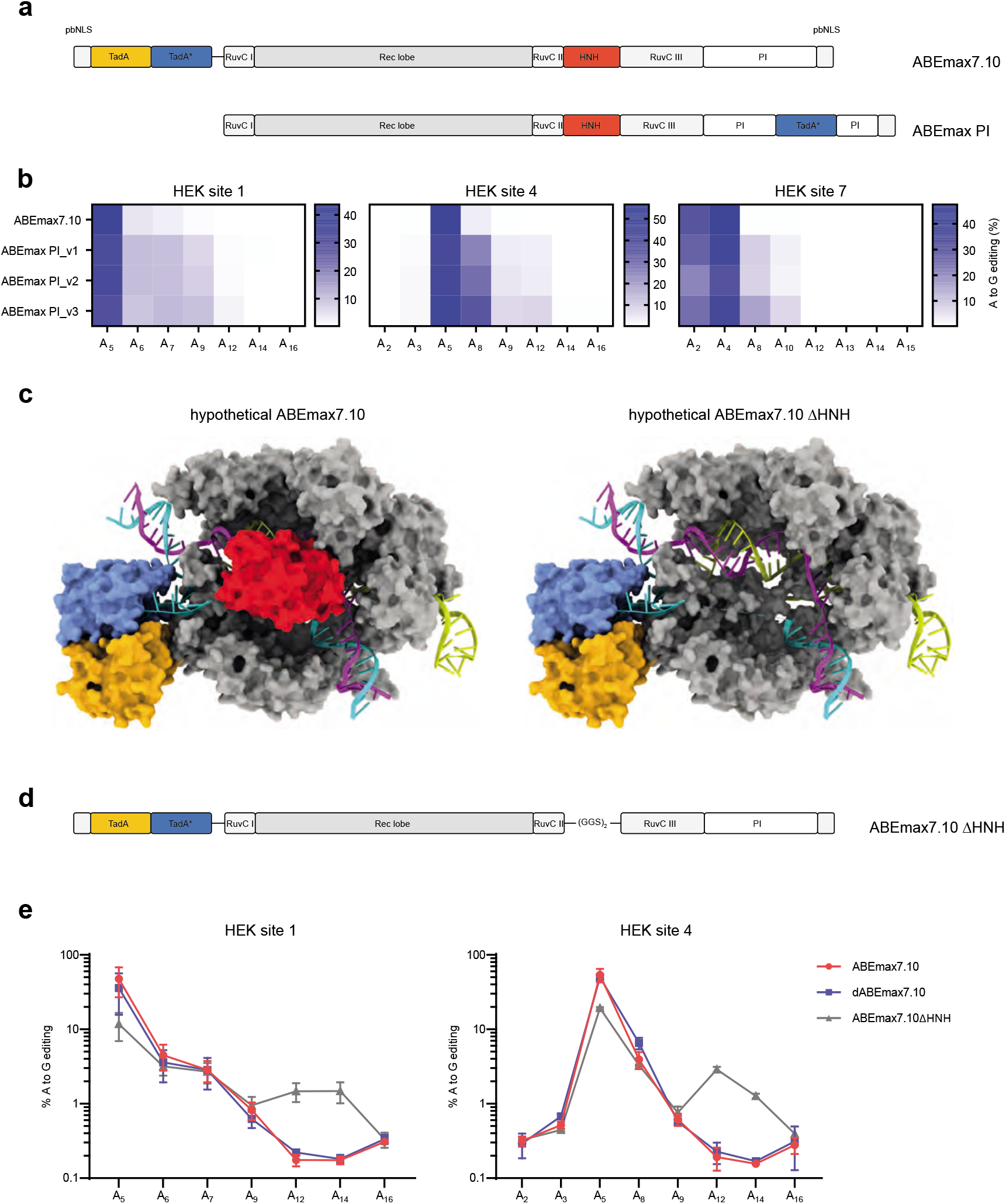
Approaches to increase substrate accessibility for base editing at PAM-proximal bases. **a)**Schematic domain organization of ABEmax7.10 and ABEmax PI1-3. ABEmax PI1-3 comprise an engineered *Sp*Cas9 (D10A) construct, where the TadA deaminase is integrated within the PI domain. ABEmax PI1, PI2, and PI3 use different linker lengths flanking the TadA deaminase. **b)**Editing efficiencies of ABEmax PI constructs are quantified by high throughput sequencing at 3 different endogenous loci. Values represent mean of three independent biological replicates performed on separate days ± s.d. **c)**Structural data (PDB: 6VPC) of adenine base editors with (left) and without (right) an HNH domain (indicated in red). Laboratory evolved, monomeric TadA deaminases indicated in orange and blue dimerize, and target the ssDNA substrate (turquoise). **d)**Schematic domain organization of ABEmax7.10 ΔHNH, where the HNH domain is replaced by a glycine-serine linker (Gly-Gly-Ser-Gly-Gly-Ser). **e)**High throughput sequencing data compares editing efficiencies of ABEmax7.10, dABEmax7.10 and ABEmax7.10 ΔHNH in endogenous loci in HEK293T cells. ABEmax7.10 retains nickase activity, nicking the target strand. Elimination of the nickase activity in ABEmax7.10 by introducing a H840A mutation in the HNH domain leads to dABEmax7.10. Complete removal of the HNH domain (that also abolishes nickase activity) in ABEmax7.10 results in ABEmax7.10 ΔHNH. Values represent mean of three independent biological replicates performed on separate days ± s.d.

Here, we report a novel ABE variant, where the HNH nuclease domain of *Sp*Cas9 is replaced by TadA. Compared to classical ABEs this variant has a PAM-proximally shifted editing window, and is significantly reduced in size.

## RESULTS AND DISCUSSION

Structural data suggest that the HNH nuclease domain (amino acids 775-908) in *Sp*Cas9 likely prevents the deaminase from accessing the ssDNA substrate at positions 13-15 counting from the PAM-distal end of the protospacer (Fig. 1c). Importantly, the HNH domain is not required for Cas9 to bind its target DNA and form an R-loop (17), and nickase activity is not essential for base editing (Fig 1e) (17). We therefore reasoned that omission of the HNH domain might improve accessibility and editing at PAM-proximal positions. To test our hypothesis, we engineered ABEmax7.10 ΔHNH. Compared to the original ABE7.10 construct, where a heterodimer of wild-type (WT) and evolved TadA-7.10 is C-terminally linked to a “nickase” or “dead” version of *Sp*Cas9, in the ABEmax7.10 ΔHNH construct the TadA heterodimer is C-terminally fused to a *Sp*Cas9 variant that lacks the HNH domain (Fig.1d). Although the highest editing rates remained at positions that are also efficiently targeted with ABEmax7.10, ABEmax7.10 ΔHNH allowed editing at positions 12 and 14 (Fig. 1e). Our data therefore suggest that the HNH domain of *Sp*Cas9 indeed constrains access of TadA to PAM-proximal ssDNA nucleobases.

Simply expanding the editing window of BEs has the disadvantage that it can generate additional undesired non-target nucleotide (bystander) edits within the ssDNA of the R-loop. This led us to investigate whether directly replacing the HNH domain with the adenine deaminase (HNHx-ABE) leads to a shift rather than a broadening of the editing window (Fig. 2a, b). Before incorporating a deaminase domain in place of the HNH domain in *Sp*Cas9, we first exchanged the HNH domain for a superfolder (sf)GFP. Transfection of these constructs into HEK293T cells demonstrated localization of GFP to the nucleus (Fig. 2c), indicating that the HNH domain can be exchanged with another protein domain without substantial misfolding and degradation of Cas9. In a next step, we engineered HNHx-ABEmax7.10 by incorporating the evolved deaminase domain from ABEmax7.10 (4) with 20 different peptide linker combinations into *Sp*Cas9 lacking the HNH domain. Using high throughput sequencing (HTS), editing efficiencies of constructs with different linker combinations were compared two days after transfection (Fig. 2d). In the most promising construct, a GGS-linker was used to join *Sp*Cas9 S793 to the TadA7.10 N-terminus, and a SGG-linker was used to join the TadA7.10 C-terminus to *Sp*Cas9 R919. This construct (HNHx-ABEmax7.10) exhibited a distinct shift in the editing window PAM-proximally, with up to 13.7% editing five days after transfection (Fig. 3a). Notably, when we exchanged the HNH domain with the cytosine deaminases FERNY, a laboratory evolved APOBEC variant (19), we observed a similar shift in the editing window, albeit with very low editing rates (Suppl. Fig. 1). While these could be increased by exchanging FERNY with the cytidine deaminase PmCDA1, HNHx-PmCDA1 constructs induced broad deamination across the entire protospacer region, leading to significant bystander edits (Suppl. Fig. 1).

**Fig 2.**
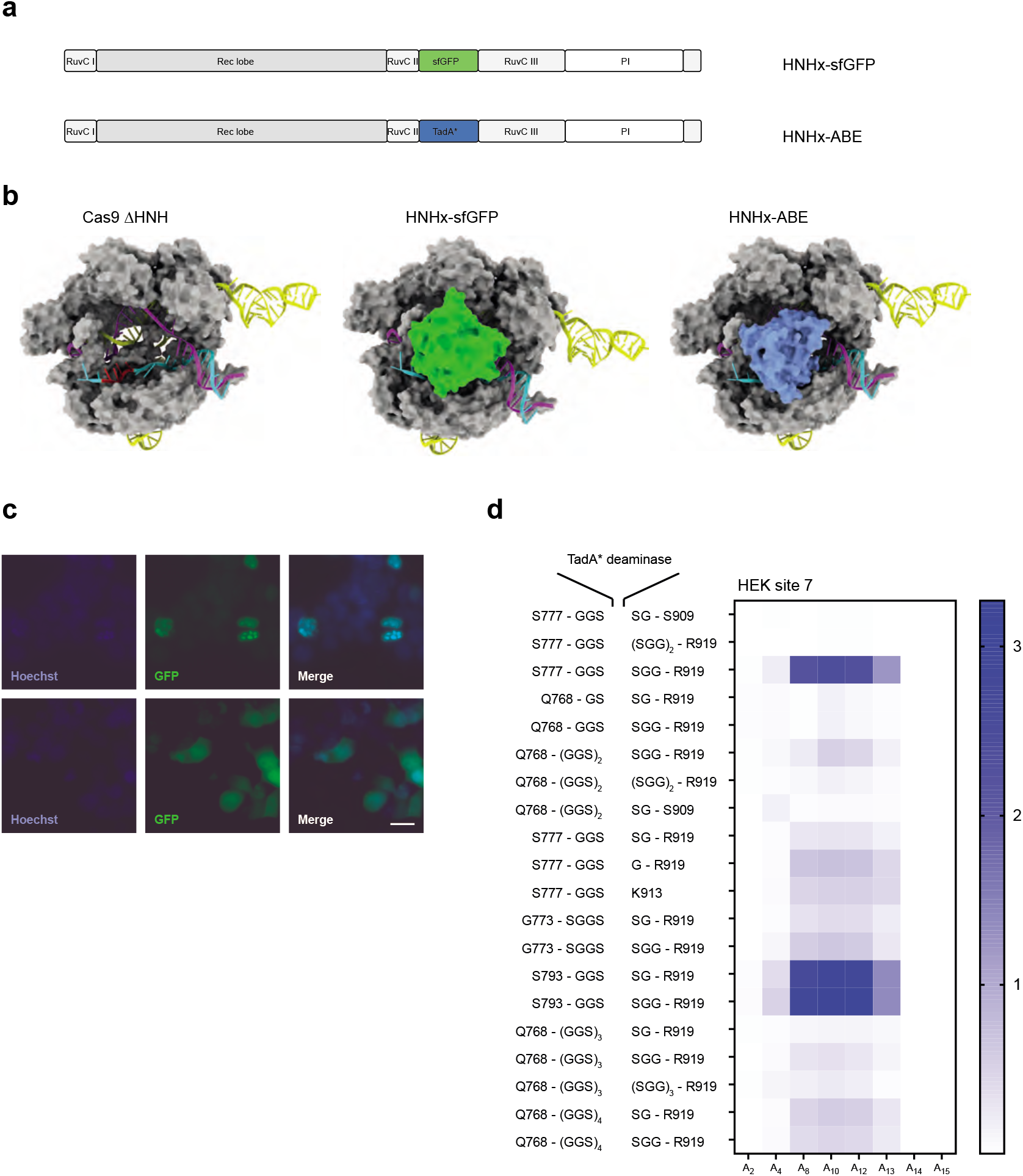
HNH domain substitution with sfGFP and ssDNA-specific deaminase domains. **a)**Schematic domain organization of Cas9 variants, where the HNH domain is replaced by a sfGFP or a ssDNA-specific deaminase domain. **b)**Structural data of hypothetical Cas9 (PDB: 5F9R) constructs, where the HNH domain is omitted, replaced by sfGFP (PDB: 2B3P) or a TadA deaminase (PDB: 6VPC). Nucleotides (14–17) indicated in red are outside of a typical editing window of ABEmax7.10 **c)**Fluorescence microscopy of HEK293T expressing Cas9, where the HNH domain is replaced with sfGFP, with (top panel) and without (bottom panel) nuclear localization signals. Blue: Hoechst, green: GFP, scale bar: 20 μm. **d)**Heat map depicting different flanking amino acids of Cas9 and linkers to incorporate the TadA deaminase in place of the HNH domain. The TadA deaminase reading frame is as listed in the Supplementary Information. Editing efficiencies were quantified by high throughput sequencing indicated as % A-to-G conversion 2 days after transfection. The x-axis defines base conversions at adenine base positions within the protospacer.

**Fig 3.**
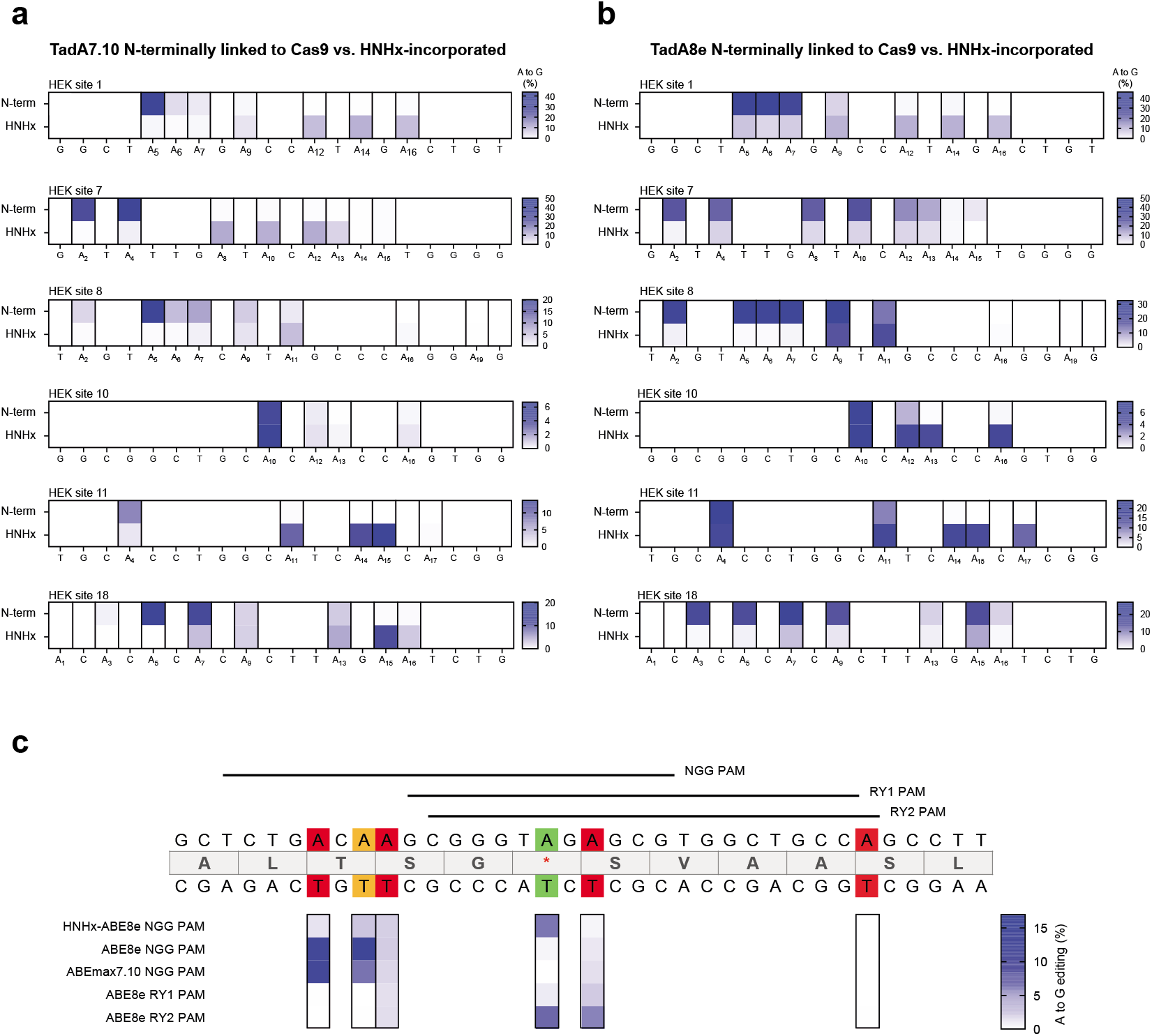
Targeting of endogenous loci. **a)**Editing efficiencies of different adenine bases within the protospacer region comparing ABEmax7.10. and HNHx-ABEmax7.10 in endogenous loci in HEK293T cells. Numbering starts with PAM-distal nucleotides. Values represent mean of three independent biological replicates performed on separate days ± s.d. **b)**Editing efficiencies of different adenine bases within the protospacer region comparing ABE8e and HNHx-ABE8e in endogenous loci in HEK293T cells. Numbering starts with PAM-distal nucleotides. Values represent mean of three independent biological replicates performed on separate days ± s.d. **c)**Editing of a disease-causing c.3188G>A mutation in the FANCA gene comparing different ABEmax7.10, ABE8e and HNHx-ABE8e constructs. Green indicates the target base, orange a synonymous mutation and red a non-synonymous mutation. Values represent mean of three independent biological replicates performed on separate days ± s.d.

Phage assisted non-continuous and continuous evolution has enabled the development of TadA variants with increased deaminase activity (20). To assess whether the use of these variants would improve editing rates of HNHx-ABEs, we next exchanged the TadA7.10 deaminase domain with hyperactive TadA8e, resulting in HNHx-ABE8e. As expected, we observed higher editing rates with HNHx-ABE8e compared to HNHx-ABEmax7.10 (up to 19.7% - Fig.3b). Higher editing efficiencies, however, came at the cost of lower specificity, and the editing window was significantly broadened (Fig 3b). Depending on the target locus, base editing with HNHx-ABE8e could therefore enhance the chances of generating deleterious bystander mutations.

Several recent studies have shown that ABEs induce substantial sgRNA-independent off-target deamination on the transcriptome, raising concerns about potential risks for clinical application (21–23). Considering that TadA is not terminally fused but instead integrated into Cas9, and thus potentially constrained by HNHx-ABE architecture (Fig. 1b and 2c), we reasoned that transcriptome-wide off-target editing might be reduced. However, while RNA-sequencing showed a decrease in off-target adenine deamination in HNHx-ABE8e treated cells compared to ABE8e treated cells, HNHx-ABEmax7.10 showed a slight increase in off-target adenine deamination compared to ABEmax7.10 (Suppl. Fig. 2). Interestingly, Gaudelli et al. recently reported an elevation in RNA off-target deamination when a TadA7.10 was fused to Cas9 as a monomer instead of a heterodimer (24), suggesting that enhanced transcriptome deamination in HNHx-ABEmax7.10 could result from the inability of TadA7.10 to form a heterodimer with wild-type TadA.

To provide an example for the potential benefit of HNHx-ABE constructs over ABEs with N-terminally fused TadA, we next targeted a disease-causing mutation in the Fanconi anaemia complementation group A (FANCA) gene (c.3188G>A - Fig. 3c). Here, the only available canonical NGG PAM site positions the disease-causing mutation outside of the editing window of classical ABEs. When we transfected cells containing the FANCA c.3188G>A mutation with HNHx-ABE8e, we obtained considerably higher on-target editing compared to ABE8e or ABEmax7.10 (Fig. 3c). Notably, similar on-target editing could be obtained when using ABE8e-SpRY^14^, where an evolved Cas9 binds a downstream NCC PAM site, positioning the disease-causing mutation within reach of classical ABEs (Fig. 3b). However, compared to HNHx-ABE8e, ABE8e-SpRY led to more bystander mutations that resulted in non-synonymous mutations in the target gene. For correcting the FANCA c.3188G>A mutation, HNHx-ABE8e therefore provides a valuable alternative to current ABE variants.

In this work we demonstrate that replacing the HNH domain of *SpCas9* with a deaminase domain shifts the editing window of BEs PAM-proximally (from positions 4-8 to 9-16 for TadA7.10). This expands the targeting scope of currently available base editors, and enables targeting of additional disease-causing mutations. Replacement of the HNH domain with TadA furthermore reduces the size of ABEs from 5.4kb to below 4.4kb, allowing the construction of single adeno-associated virus (AAV)-vectors with minimal promoters for *in vivo* delivery. It is important to note, however, that editing rates of HNHx-ABEs were lower compared to ABEs with N-terminally fused TadA. In part, this is due to the absence of an HNH domain, as ABEmax7.10 lacking the HNH domain (ABEmax7.10 ΔHNH) also exhibited a reduction in editing rates compared to classical ABEmax7.10. Interestingly, this difference in editing was not caused by the lack of target strand nicking (dABEmax7.10 lacking nickase activity performed comparably to ABEmax7.10), suggesting that the role of the HNH domain in base editing goes beyond nicking of the target strand, and that it potentially also contributes to the structural integrity of the BE complex. Another potential reason for lower editing rates of HNHx-ABE constructs could be steric hindrance by the surrounding Cas9 protein and prevention of TadA dimerization (Suppl. Fig 3), as recent studies have demonstrated that TadA typically forms a dimer in its functional state (4, 20, 25). In the future, directed protein evolution or rational protein engineering strategies may nevertheless further refine HNHx-ABE constructs and enable editing efficiencies similar to classical ABEs. In addition, our approach of replacing the HNH domain could be extended to other effector enzymes that act on ssDNA, further expanding the Cas9-based genome editing tool box.

## DATA AVAILIBILITY

All data will be made available upon reasonable request. Amino acid sequences of BE constructs and primer sequences for the sgRNAs are shown in Supplementary data file 1.

## ACCESSION NUMBERS

HTS data from all experiments will be deposited online before publication.

## ACKNOWLEDGMENTS

We would like to thank Weihong Qi of the Functional Genomics Center Zurich for the analysis of RNA-seq experiments. This work was supported by the Swiss National Science Foundation grant (#180257), and by the PHRT grant (#528).

## AUTHOR CONTRIBUTIONS

L.V. designed the study, performed experiments, analysed data, and wrote the manuscript. L.S., K.M., T.R. and N.M. performed experiments and analysed data. G.S. designed and supervised the research and wrote the manuscript. All authors approved the final version.

---

**Suppl. Fig S1.**
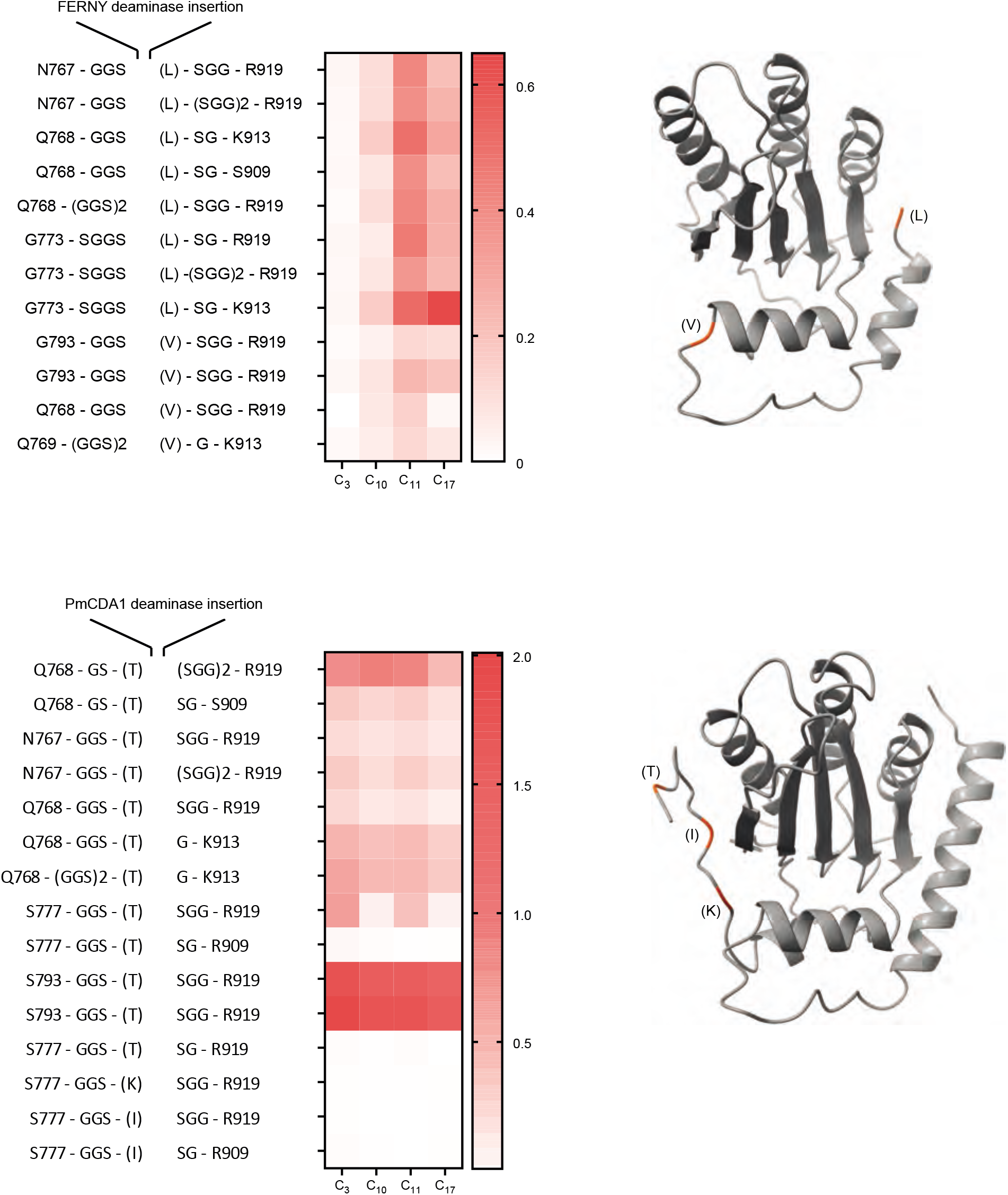
Heat map depicting different flanking amino acids of Cas9 and linkers to incorporate the FERNY (top) and PmCDA1 (bottom) deaminases in place of the HNH domain. The FERNY and PmCDA1 deaminase reading frames are listed in the Supplementary Information. Linkers that join Cas9 and deaminase at different amino acid positions within the deaminase are indicated in brackets.

**Suppl. Fig S2.**
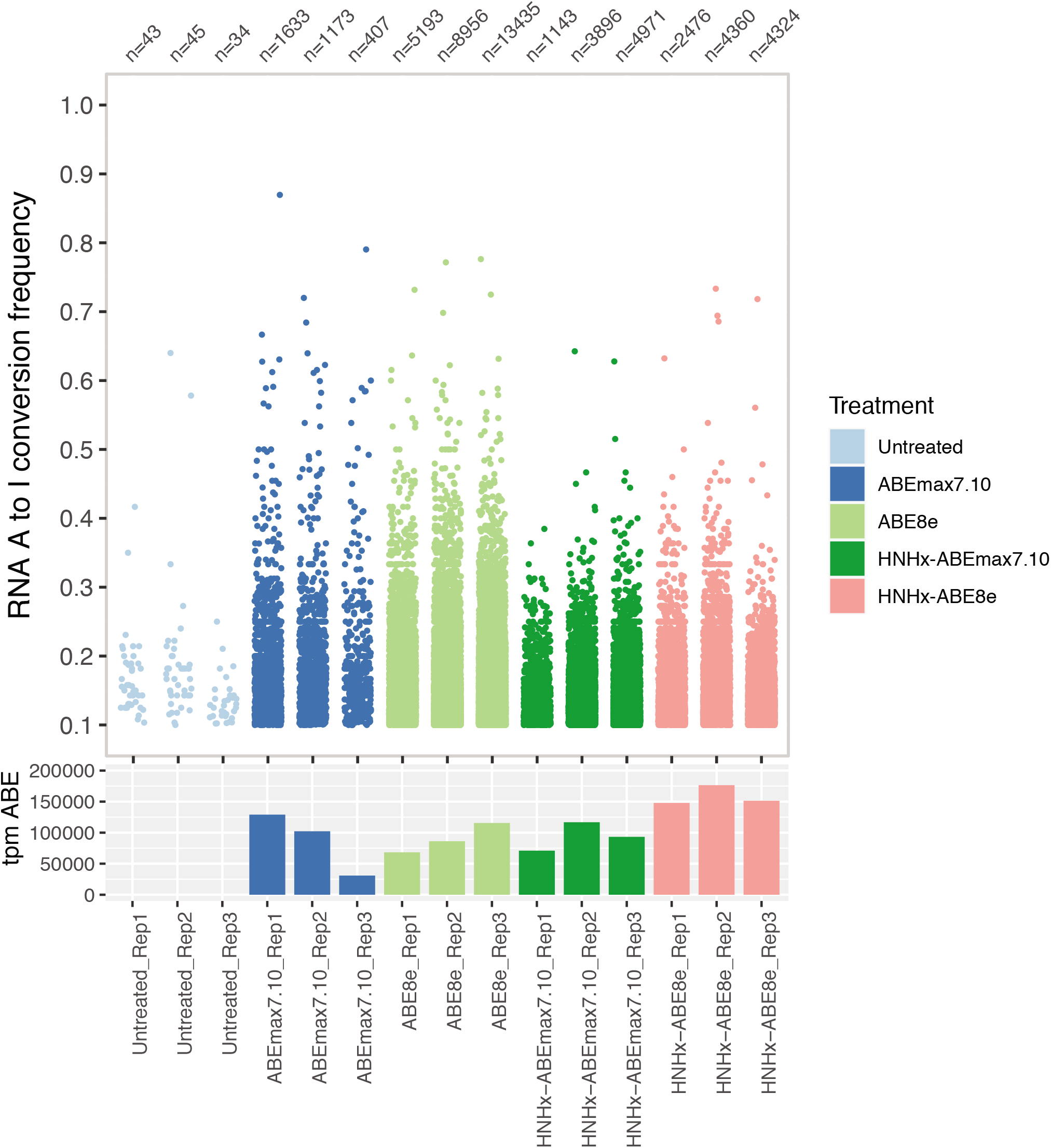
RNA-seq analysis determines transcriptome-wide RNA off-targets after transfection of ABEmax7.10, ABE8e, HNHx-ABEmax7.10 and HNHx-ABE8e constructs.

**Suppl. Fig S3.**
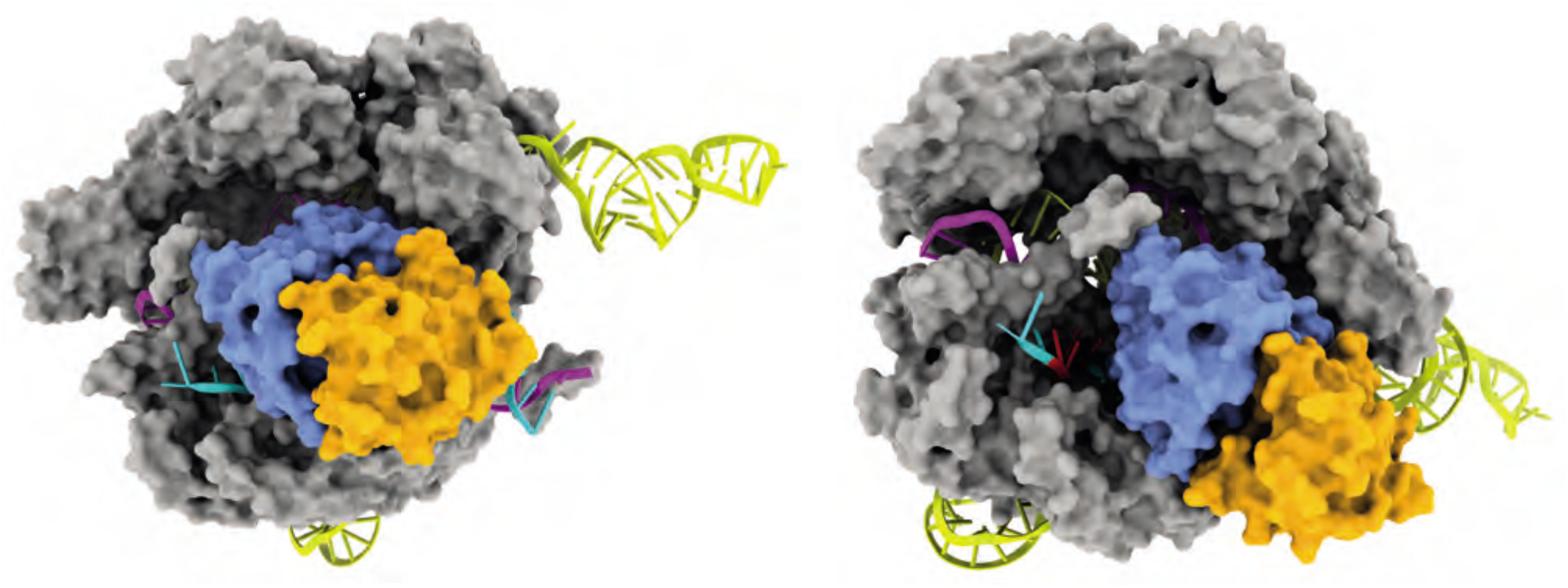
Different viewing angles of a hypothetical HNHx-ABE variant with a dimerized TadA. Subunits are colored as follows: Cas9 ΔHNH, grey; TadA linked to Cas9, blue; TadA from another molecule dimerizes to Cas9-bound TadA for functional deamination, orange.

## METHODS

### General methods and cloning

PCR was performed using Q5 High-Fidelity DNA Polymerase (New England Biolabs). All base editor constructs were assembled using NEBuilder HiFi DNA Assembly (New England Biolabs). Plasmids expressing sgRNAs were cloned using T4 DNA Ligase (New England Biolabs).

### Cell culture and high-throughput sequencing

HEK293T cells (ATCC CRL-3216) were cultured in Dulbecco’s modified Eagle’s medium GlutaMax (Thermo Fisher Scientific), supplemented with 10% (v/v) fetal bovine serum (FBS) and 1% penicillin-streptomycin (Thermo Fisher Scientific) at 37°C and 5% CO_2_. Cells were maintained at confluency below 90% and seeded on 96-well cell culture plates (Greiner). 12-16h after seeding, at approximately 70% confluency, cells were transfected using 0.5μl Lipofectamine 2000 (Thermo Fisher Scientific) and 400ng base editor plasmid DNA and 100ng sgRNA plasmids. Cells were incubated for 5 days. Genomic DNA was isolated by adding 10μl lysis buffer (10mM Tris-HCl at pH8.0, 2% Triton X and 1mM EDTA and 25μg/ml Proteinase K) to 30μl cell suspension. The lysate was incubated at 60°C for 60min, followed by a 95°C incubation for 10min. The lysate was diluted with ddH2O to a final volume of 100μl. 2μl of the diluted lysate was used for subsequent PCR reactions of 10μl using NEBNext High-Fidelity 2x PCR Master Mix. The PCR product was purified using Agencourt AMPure XP beads (Beckman Coulter), and amplified with primers containing sequencing adapters. The products were gel purified and quantified using the Qubit 3.0 fluorometer with the dsDNA HS assay kit (Thermo Fisher Scientific). Samples were sequenced on an Illumina MiSeq.

### HTS data analysis

Sequencing reads were demultiplexed using MiSeq Reporter (Illumina), and analysed using a Matlab script as previously described (5). Values are shown as n=3 independent biological replicates over different days, with mean ± s.d.

### Microscopy

HEK293T cells were transfected with 50ng GFP-expressing plasmids in a 96 well plate, counterstained with Hoechst 33342 and imaged using a Zeiss Apotome. Imaging conditions and intensity scales were matched for all images. Images were analysed using Fiji ImageJ software (v1.51n).

### Linker determination and testing

Structural data from *Sp*Cas9 (PDB: 5F9R) was used to estimate linker lengths flanking deaminases in PyMol version 2.3.4. Different constructs with combinations of N- and C-terminal linkers were tested, editing efficiencies and activity windows determined by high-throughput sequencing.

Molecular graphics and analyses were done with UCSF ChimeraX, developed by the Resource for Biocomputing, Visualization, and Informatics at the University of California, San Francisco, with support from National Institutes of Health R01-GM129325 and the Office of Cyber Infrastructure and Computational Biology, National Institute of Allergy and Infectious Diseases.

### RNA-seq experiments and data analysis

RNA library preparation was performed using the TruSeq Stranded Total RNA kit (Illumina) with a ribosomal RNA (rRNA) deletion step. RNA-seq libraries were sequenced on an Illumina Novaseq machine with a sequencing depth of 140-200 Mio reads per sample.

### Quality control, pre-processing and alignment of RNA-seq reads

Quality of Illumina PE RNA-seq reads was evaluated by FastQC v0.11.7 (https://www.bioinformatics.babraham.ac.uk/projects/fastqc/). Possible contaminations (genomic DNA, rRNA, Mycoplasma, etc.) were screened for using FastqScreen v0.11.1 (https://www.bioinformatics.babraham.ac.uk/projects/fastq_screen/) against a customized database that consists of SILVA rRNA (https://www.arb-silva.de/), UniVec (https://www.ncbi.nlm.nih.gov/tools/vecscreen/univec/), refseq mRNA sequences and selected genome sequences (human, mouse, arabidopsis, bacteria, virus, phix, lambda, and mycoplasma) (https://www.ncbi.nlm.nih.gov/refseq/). Illumina PE reads were pre-processed using Trimmomatic version 0.36 to trim off sequencing adaptors and low quality ends (average quality lower than 20 within a 4 nt window). Flexbar version 3.0.3 was used to remove the first 6 bases of each read, which showed priming bias introduced during library preparation. PE RNA-seq reads were generated with different read length (2X51 and 2X151). After adapter and quality trimming, if the read length was longer than 50 nt, only the first 50 nt were kept for downstream STAR mapping and variant calling. Quality controlled reads (average quality 20 and above, read length 20 and above) were aligned to the reference genomes (human reference genome: GRCh38.p10, Ensembl release 91) using STAR version 2.7.0e with 2-passes mode. PCR-duplicates were marked using Picard version 2.9.0. Read alignments were comprehensively evaluated in terms of different aspects of RNA-seq experiments, such as sequence quality, gDNA and rRNA contamination, GC/PCR/sequence bias, sequencing depth, strand specificity, coverage uniformity and read distribution over the genome annotation, using R scripts in ezRun (https://github.com/uzh/ezRun/) developed at the Functional Genomics Center Zurich.

### RNA sequence variant calling and filtering

Variant calling from RNA-seq reads was performed according to GATK Best Practices (https://gatkforums.broadinstitute.org/gatk/discussion/3891/calling-variants-in-rnaseq). In detail, GATK (version 4.1.2.0) tool SplitNCigarReads was applied to postprocessed the read alignments. Afterwards variants were called using HaplotypeCaller (GATK version 4.1.2.0) on PCR-deduplicated, post-processed aligned reads. Variant loci in base-editor overexpression experiments were filtered to exclude sites without high-confidence reference genotype calls in the control experiment. For a given SNV, the read coverage in the control experiment should be >90th percentile of the read coverage across all SNVs in the corresponding overexpression experiment. Only loci having at least 99% of reads containing the reference allele in the control experiment were kept. Only sites with more than 10 reads mapping in the overexpression experiment were kept. The cutoff for RNA A to I conversion frequency was set to 0.1.

